# *Drosophila* Video-assisted Activity Monitor (DrosoVAM): a versatile method for behavior monitoring

**DOI:** 10.1101/2025.07.29.667363

**Authors:** Maxime Revel, Emi Nagoshi, Robert K Maeda

**Affiliations:** University of Geneva

## Abstract

*Drosophila melanogaster* has been a pioneering model system for investigations into the genetic bases of behavior. Studies of circadian activity were some of the first behaviors investigated in flies. The DAM system by TriKinetics played a key role in establishing the fundamental feedback loop of the circadian clock. Although this method has many time proven to be extremely useful, it suffers from its simplification of activity to the interruption of an infrared beam. It is blind to fly movements not disrupting the beam and any modifications to this assay to achieve better resolution often requires the purchase of new and expensive modules. We required a relatively high-throughput system to explore the potential post-mating activity changes of larger *Drosophila* species. Rather than investing in a larger and more-complex DAM system, we designed a new monitoring system that is more versatile, economic and sensitive than DAM. This new system, called DrosoVAM (Drosophila Video-assisted Activity Monitoring), is simple to implement and cost efficient, using a Raspberry Pi-controlled infrared, digital video system to record multiple chambers and Python scripts that drives the deep learning software DeepLabCut, to track fly activity over multiple days.

## Introduction

Model organisms continue to play a critical role in deciphering the links between neuronal circuits and complex behaviors (Brenman‐Suttner et al., 2020; Cooper and Raamsdonk, 2018; Martín and Alcorta, 2017; Orger and Polavieja, 2015; White, 2016). In particular, *Drosophila melanogaster* has helped in unravelling the cellular and molecular basis of complex behaviors like memory formation (Busto et al., 2015; Tully et al., 1994), aggression (Gaspar et al., 2022; Zwarts et al., 2012), sleep (Dubowy and Sehgal, 2017) and courtship (Philipsborn et al., 2011; Raun et al., 2021; Yamamoto and Koganezawa, 2013). Recently, the fruit fly has even proven to be useful in modeling behaviors found in human neurodegenerative diseases such as Alzheimer’s disease (Tsuda and Lim, 2018) and Parkinson’s disease (Nagoshi, 2018).

As our understanding of *Drosophila* behavior grows, we are becoming increasingly aware of the limitations of some of the standardized assays. One frequently used example of this is the TriKinetics DAM system (Chiu et al., 2010) used to monitor locomotor activity. This method places individual flies into narrow tubes with a minimal food source (5% Sucrose, 2% Agar) located at one end. Multiple tubes are then inserted into a monitor that emits an infrared (IR) beam through the center of each tube and records the number of times the fly disrupts the IR beam. Due to its simplicity of use and high throughput nature, this system has proven to be extremely useful. However, its spatial resolution is low, as the fly movements are generally monitored only in the center of the tube. Furthermore, the restricted nature of the chambers limits the movements of the fly in unnatural ways.

Due to the limitations of methods like DAM, different approaches are being developed to track the behavior of individual flies more precisely (Sitaraman et al., 2024). Some of these systems use deeplearning algorithms to track, define and quantify micro-movements such as proboscis extension, or repositioning of halteres and antennae (Keleş et al., 2025). Other software and protocols are more specifically designed to identify behaviors of interest such as egg-laying (Bräcker et al., 2019) or feeding (Niu et al., 2021). In contrast to DAM, many of these methods surprisingly suffer from being too high resolution (Bräcker et al., 2019; Niu et al., 2021; Qu et al., 2022), making data analysis more difficult and thereby reducing the number of individuals analyzed per experiment.

In our lab, we were interested in using a high-throughput system like DAM to examine potential postmating changes in the activity of other *Drosophila* species like *D. virilis* and *D. hydei*. However, initial trials using the DAM system highlighted some of its limitations. First, the tubes used for the DAM system present in our lab (designed for *Drosophila melanogaster*) were too narrow to support the free movement of larger species like *D. virilis* and *D. hydei*. Second, the minimal food source used in DAM assays is known to be insufficient to support normal egg-laying of females (Mirth et al., 2019) and our attempts to replace the minimal food with standard, rich food proved to be problematic due to its propensity to dry out in such a small volume and the obscuring influence of hatching larvae over time.

To circumvent these difficulties, we designed a new monitoring system that is simple, more versatile, economic and sensitive than the DAM monitoring technique. Dubbed DrosoVAM for *Drosophila* Video-assisted Activity Monitor, this system is made up of three parts. The first part consists of an infrared digital video system that records flies in 3D printed chambers illuminated by an infrared light source. As the circadian clock of *Drosophila* is not altered by infrared light (Frank and Zimmerman, 1969), infrared monitoring can be used to track flies during both day and night cycles (as imposed by changing visible light) with equal sensitivity. In the second part, the digital recordings are cropped and then analyzed by the deep learning software DeepLabCut (Mathis et al., 2018; Nath et al., 2019) to track the flies in the chambers and convert the videos to positional values. And last, in the third step, a set of Python scripts are run to interpret the tracking data and infer the movement behaviors of the flies. The versatility of the python script, as well as the ease of 3D printing, makes the method highly flexible and inexpensive to modify in order to test for activity in a variety of situations. Here, we demonstrate the utility of this method in basic activity monitoring as well as in a post-mating food preference assay.

## Materials and Methods

The DrosoVAM system {Fig. 1} is designed as a three-step protocol, each step of which is described here. All software used, STL files for 3D printing, along with the hardware list, can be found on the associated GitHub repository {https://github.com/LabMaeda/DrosoVAM/}.

**Figure 1.**
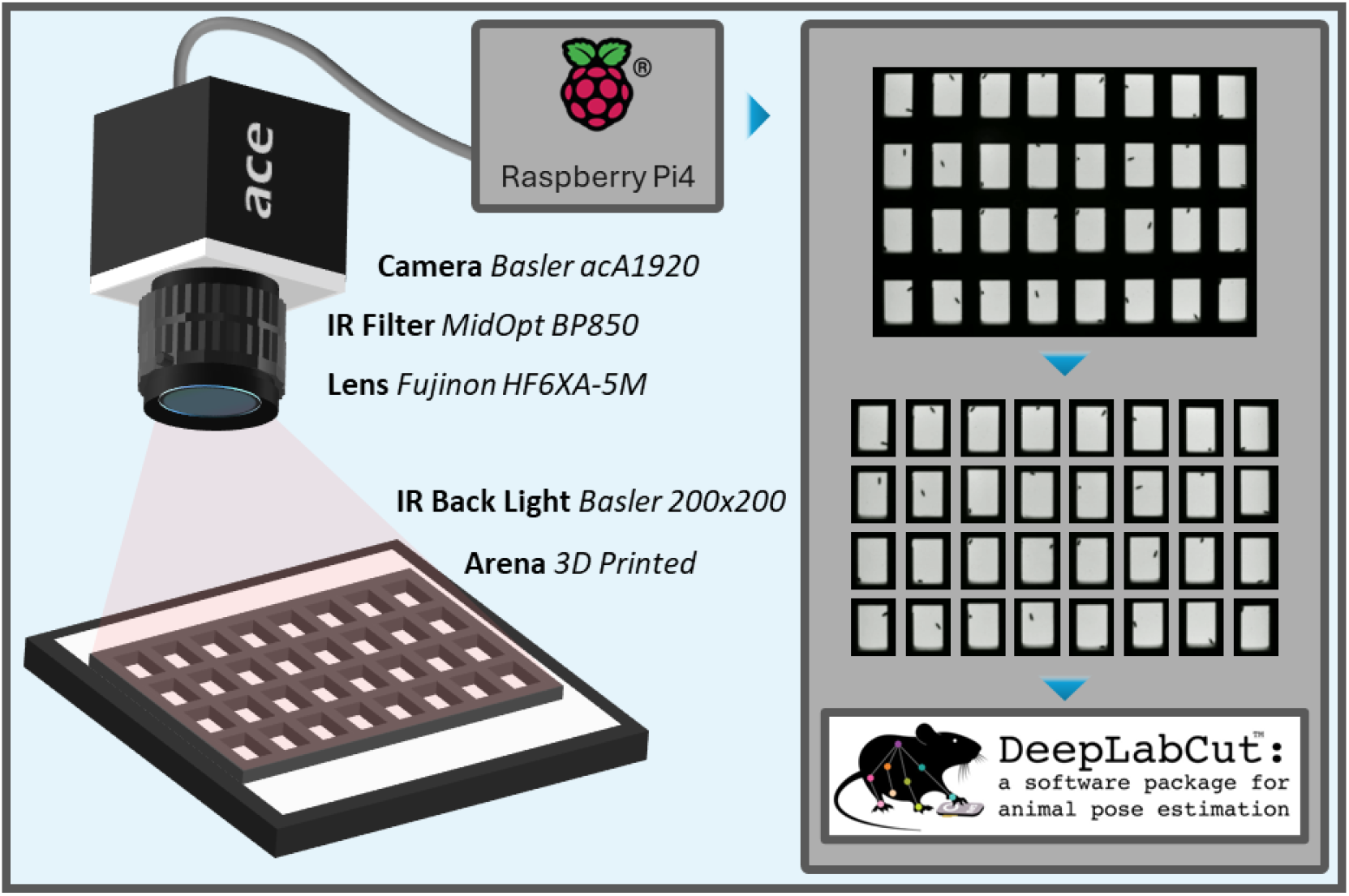
Hardware used to monitor flies using DrosoVAM and overview of the video analysis. Flies are placed in a chamber laying horizontally on an infrared backlight and are recorded from above using a camera equipped with an infrared pass filter and lens. The camera is connected to a Raspberry Pi4 on which a custom script is run to capture videos, here at 5fps to limit file size. Videos are then cropped into individual videos each focused one chamber and analyzed using the tracking software DeepLabCut.

### Hardware – Video acquisition and preparation

To ensure maximum versatility and ease of troubleshooting, we developed our own recording arena, 3D-printed in polylactic acid using a Prusa Mk3 filament printer. The recording arena measures 196×145mm and is enclosed between two removable, 2 mm-thick acrylic sheets (purchased through various suppliers on amazon.fr). The locomotor activity arena (Fig. S1a) is composed of 32 chambers (23 × 15 × 5 mm), each with a small opening (3 mm diameter), through which the fly can be inserted, and a 5mm wall at the “bottom” to help contain the food while pouring. Alternatively, a food preference arena (Fig. S1b) was designed with 24 chambers (50 × 10 × 5 mm) that have two walls to support food at each extremity.

The arena, once filled with standard cornmeal food and loaded with flies, was placed horizontally on top of an infrared light source (Basler IR Back Light 200×200). Recording was performed using a 1920×1080p camera (Basler ac1920-25um) equipped with a lens (Fujinon HF6XA-5M) and an IR-pass filter (MidOpt BP850) (Fig. 1). The filter used on the camera lens completely blocked any non-IR light, so images recorded during either light-or dark-phases are identical, and tracking is not impaired by shifting light and dark phases.

The camera was connected to a Raspberry Pi 5 running a custom Python script that uses the pypylon wrapper from Basler AG to produce series of 2h long videos segments at 5 fps. This relatively low frequency allowed the videos to not exceed 400MB, thus avoiding working with large files, all while keeping enough time resolution to monitor fly movements in the chambers. Although the recording was performed using a Raspberry Pi computer, more powerful machines could also be used if they are able to run Python scripts and to store the videos.

Videos were then cropped into smaller videos, each focusing on one chamber, using a second Python script that uses ffmpeg-python. This script works by defining the arena in a graphical interface to divide the whole viewing frame into a defined number of columns and rows based on the arena being used.

### Software I – Tracking

To optimize the tracking of the flies in the chambers, we took advantage of the DeepLabCut software (Mathis et al., 2018; Nath et al., 2019) which allows for marker-less tracking of various model animals. The string of 2-hour videos of each chamber were loaded into the software and analyzed using a trained network to identify flies at our level of resolution. The network we used, called “DrosoVAM_V3”, has been trained from a ResNet-50 network for 750,000 iterations using around one thousand labelled frames extracted from several videos. Only frames where a fly’s position was properly identifiable were labelled to avoid introduction of mislabeled data in the training.

Analyses were performed using a workstation equipped with a GPU (NVIDIA RTX-4000 Ada generation), using DeepLabCut version 2.3.9, Python version 3.10.11, TensorFlow version 2.10, CUDA version 11.8 and CudNN version 8.9.7.

### Software II – Behaviour analysis and interpretation

To transcribe tracking data into behaviors that are interpretable, we developed python scripts to read the .csv files produced by DeepLabCut, analyze them and generate plots corresponding to the information in which we were interested. To maximize the number of flies that can be monitored simultaneously, we decided to use a camera that records the chamber from a distance of around 25cm. In our experiments, this allowed for a reasonable trade-off between the number of chambers examined and tracking precision. While misdetection by DeepLabCut was rare, it occasionally happened. To help control for this, we included a step in our analysis that: 1) identifies the limits of the chamber, 2) identifies the frames where the flies are detected outside of the chamber and then 3) replaces the coordinates by the average position between the values of the preceding and next detections located inside the chamber.

For each fly, frame to frame distances were summed for the bin durations. For each 10-, 30-, or 60-minute bin, the average distance was calculated. Similarly, in the virtual-DAM, the number of detections along the midline of the chambers were summed for the bin durations, and the median of all flies was calculated for visualization. Position analyses were carried out in a similar manner, with the position of the flies being first average for each second, then averaged over the bin duration. The average position of all flies of a given mating status was then calculated for visualization. For food preference assays, the frequency of the flies localization was calculated for each fly and for each spatial bin, counting for how many frames the fly was located in each area of the chamber.

#### Drosophila Activity Monitoring (DAM) Assay

Standard Trikinetics DAM assays were carried out as described in Dorcikova et al. (Dorcikova et al., 2023) To allow mated females to lay eggs, the standard Sucrose food was replaced by standard cornmeal food.

## Results

Due to our interest in monitoring the female post-mating changes in activity in flies of different species, we sought an alternative to the traditional DAM system. For this, we developed a simple video recording-based system that we call DrosoVAM, or *Drosophila* Video-assisted Activity Monitor (detailed in Fig. 1). To confirm that DrosoVAM can track activity, we first looked at virgin female *Drosophila melanogaster* and assessed the capacity of DrosoVAM to replicate results obtained with DAM. In these experiments, we analyzed the distance covered by each virgin female in 10 minutes bins, as measured by our DrosoVAM protocol (Fig. 2a) and compared it with the number of crossings detected per 10 minutes by the DAM system (Fig. 2b). Overall, DrosoVAM largely replicates the activity pattern derived from the DAM system; the periodicity of the activity peaks surrounding the light changes is distinctly visible, particularly, the evening activity increase (Figs 2a.-b. for representative individual traces see Fig. S2). However, subtle differences can also be seen. As the measurements used by the two methods are of different natures (number of detections vs distance moved), direct statistical comparison is impossible. We attempted to bypass this by mimicking the DAM assay using our DrosoVAM videos. We did this by simulating a virtual laser in the middle of the chamber and counting how many times the fly crosses this virtual line (Fig. 2c). We then averaged several days of monitoring to compare the results of DAM with our “virtual DAM” assay (Fig. 2d). Comparing the two experiments, we find that the general pattern of activity seems to be similar, but we that the daytime activity was significantly higher when monitored using the DrosoVAM system. Furthermore, both the morning peak, triggered by light onset, and the anticipatory evening peak were more pronounced in the DrosoVAM system.

**Figure 2.**
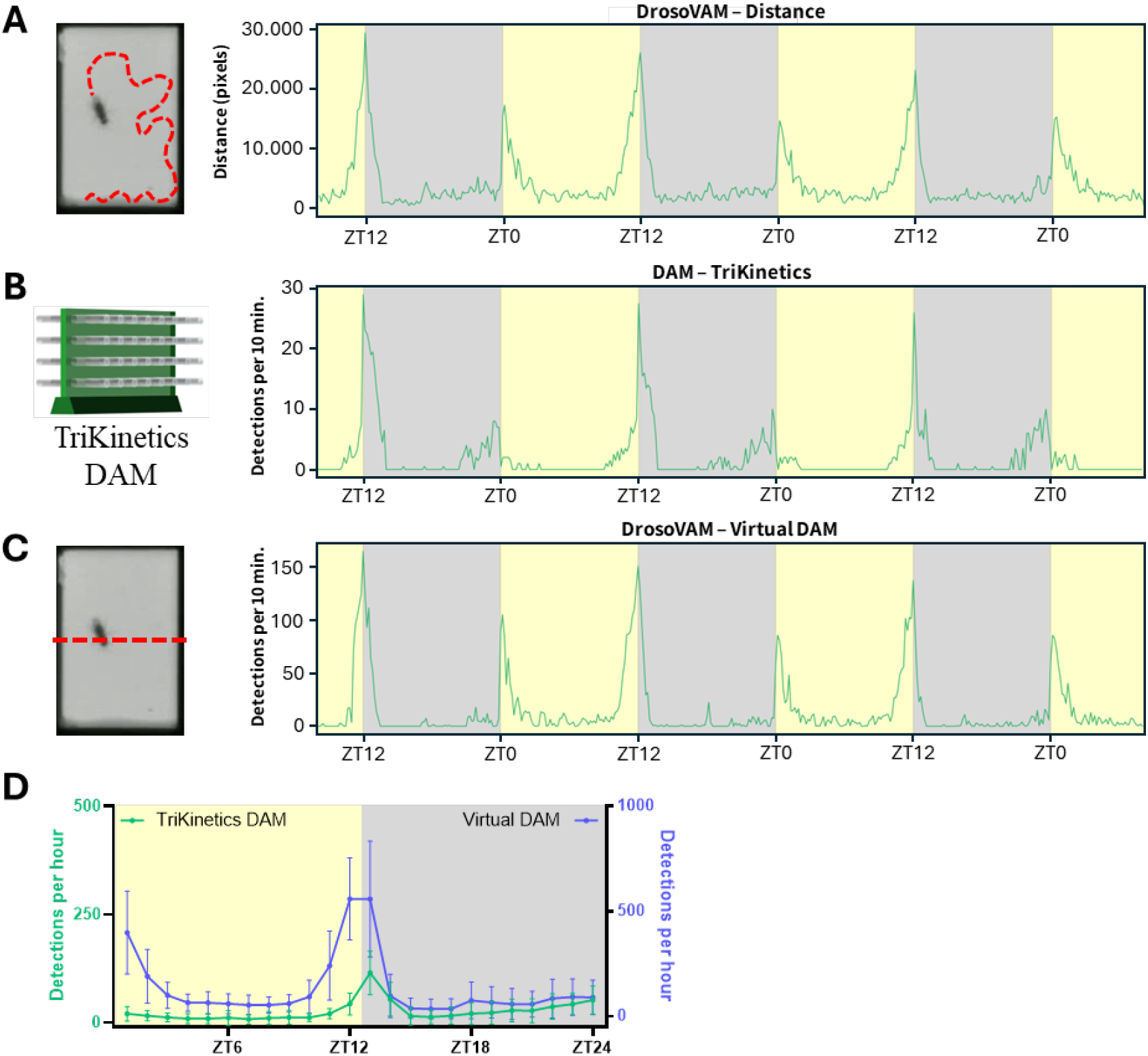
Locomotor activity of virgin *D. melanogaster*. Light/dark periods are shown with yellow/grey background respectively. A. Average distance (in pixels) moved by the flies (n= 16) estimated using DrosoVAM. B. Activity monitored using the TriKinetics DAM system (n= 32). Median number of detections per bin of 10 minutes. C. Virtual-DAM analysis simulated from DrosoVAM videos (n=16). Median number of detections per bin of 10 minutes. D. Comparison of average day activity, measured with DAM (green, left y-axis) or DrosoVAM virtual DAM (blue, right y-axis). Average day calculated from two full days of monitoring, binning of 60 minutes.

These initial results encouraged us to also examine flies in which circadian activity is disturbed. We turned to *per* null mutant flies (*per*^*01*^) that lack the endogenous molecular clock (Konopka and Benzer, 1971). Under light/dark (LD) cycles, *per*^*01*^ flies display distinct startle activity peaks at the lights-on and lights-off transitions and become arrhythmic in constant darkness (DD) (Tomioka et al., 1998).

Consistent with these previous findings, locomotor activity of *per*^*01*^ flies recorded using DrosoVAM lost all rhythmicity when switching from LD to DD (Fig. 3a), in a fashion similar to our recordings using the DAM system (Fig. 3b). Aside from this observation, we also confirmed the capacity of our system to be used for long-running experiments (here for 8 days). In our hands, we have found out that experiments can be run for up to 10 days before the food becomes too dry for flies to survive.

**Figure 3.**
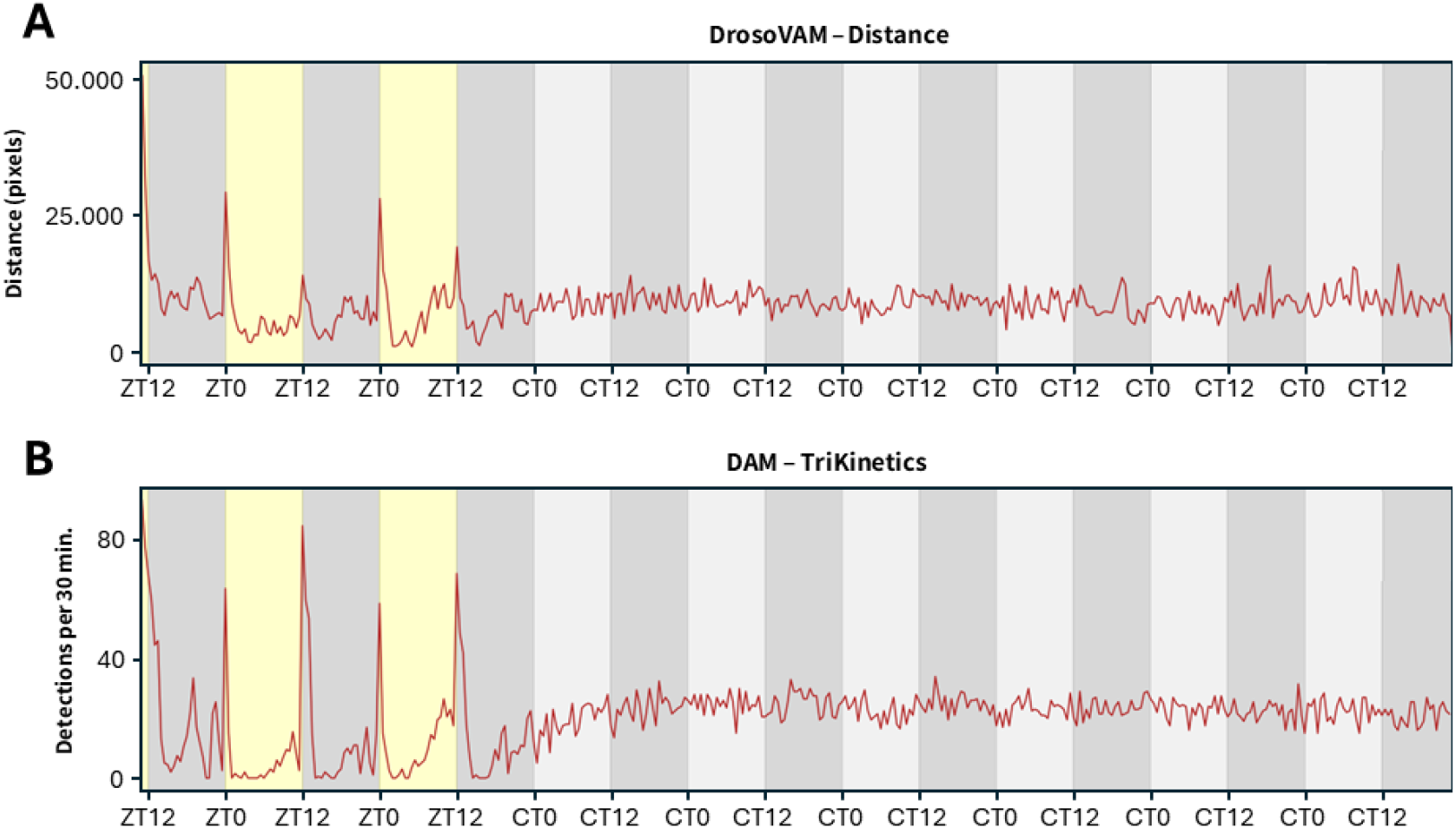
Locomotor activity of male *per*^*01*^ *D. melanogaster* recorded using the DrosoVAM and the DAM systems. Light and dark periods are shown in yellow and grey background, respectively. In constant darkness, subjective day is shown in light grey. 30 minutes bins, 204h of monitoring. A. Average distance (in pixels) moved by the flies (n=16) estimated using DrosoVAM. B. Activity monitored using the TriKinetics DAM system (n=32). Median number of detections per bin of 30 minutes.

In a first attempt at identifying behavioral post-mating responses using DrosoVAM, we placed freshly mated *Drosophila melanogaster* females into monitoring chambers for video recording over the subsequent three and a half days under a LD cycle (Fig 4a). Using this assay, we find that mated females are significantly more active during the night periods relative to virgin flies (Fig. 4b). As DrosoVAM works through video, we were able to visually confirm that these changes in activity were real and not an artefact of our tracking method. On closer examination, mated flies seem to show lower activity peaks at the light/dark transitions (Fig. 4c) combined with a reduction of the mid-day siesta and night-time sleep behaviors that characterize the circadian behavior of virgin females (fig. 4c). Using DAM, we also see a reduction in the light/dark transition activity peaks (Fig. 4d), but we do not see the overall activity changes that we see with DrosoVAM.

**Figure 4.**
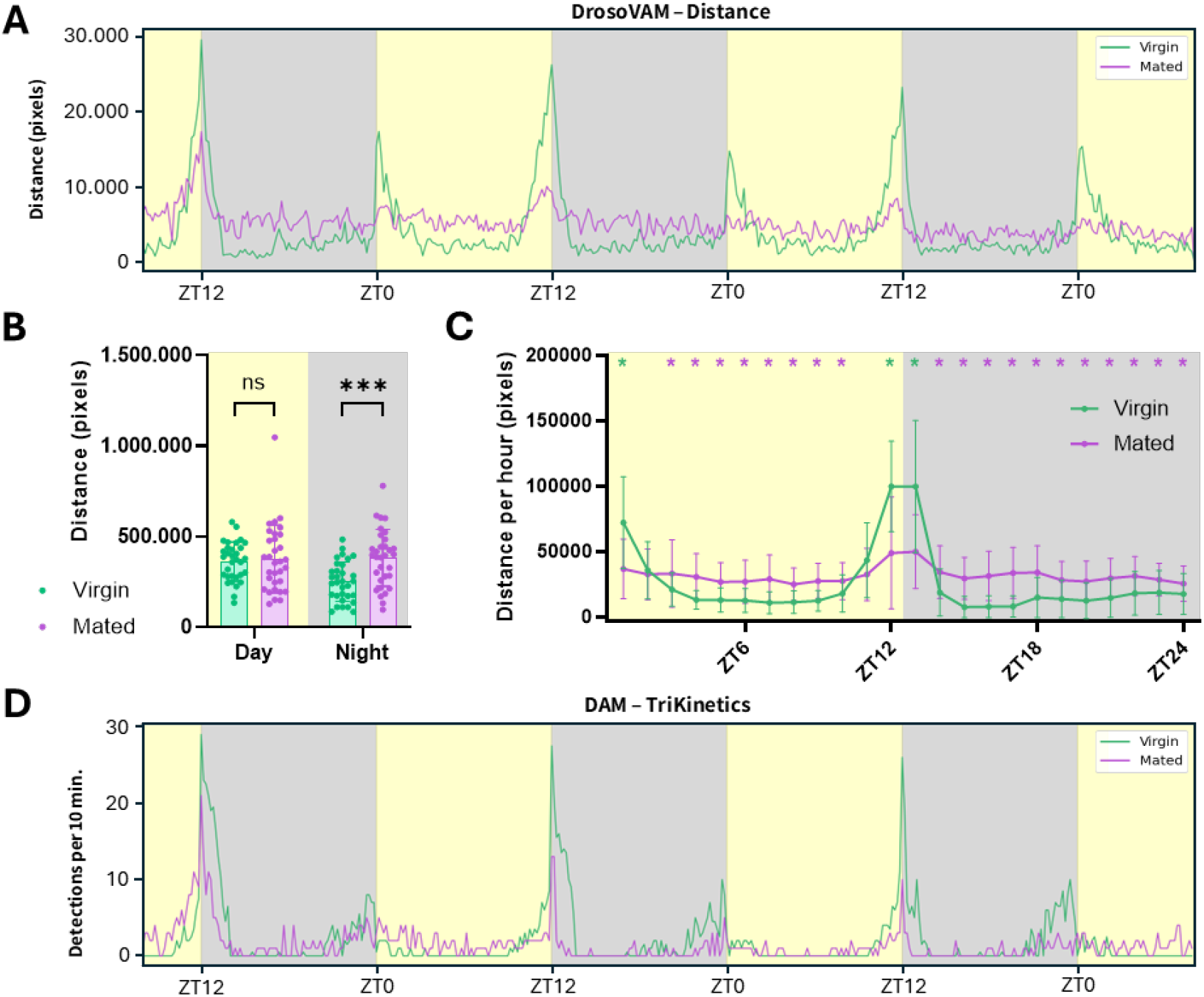
Activity comparison between *D. melanogaster* virgin and mated females. Virgins are shown in green, mated females in purple. Light/Dark periods are shown with yellow/grey backgrounds, respectively, in 10 minutes bins over 72h of monitoring. A. Average distance (in pixels) moved by the flies (mated n=16/ virgin n=16) estimated using DrosoVAM. B. total distance moved by flies during complete day and night periods (12h/12h). Mating status is compared for both periods using t-test (Day: p-value = 0.7129, Night: p-value = 0.0002). C. Comparison of the average activity measured using DrosoVAM distance analysis on virgin and mated females over a 24-hour period. The average day was calculated from two full days of monitoring, with bins of 60 minutes. The impact of mating status has been tested for each hour using multiple Mann-Whitney tests (No star means no statistical discovery, one star indicates a discovery, the color of the star indicates which one of the conditions is higher) D. Activity analysis using the TriKinetics DAM system (n=32/32). Median number of detections per bin of 10 minutes.

We wondered what could account for the differences between these two systems in both virgin and mated flies. Because our system continuously records the fly position, we mapped the fly position, overall and over time. From this analysis, we find that mated flies tend to spend most of their time moving near/on the food while virgin flies tend to stay away from the food (Fig. 5a-b). This greater proximity to the food could be explained by the need of mated females to lay their eggs on the food, along with the necessity for higher nutrient intake after mating. As smaller movement near the food would not be recorded by DAM, this behavior would go largely unnoticed. Furthermore, previous results using DAM have shown that virgin females show a higher activity in the late hours of the night relative to mated females (Riva et al., 2022). We also observe this in the hours just prior to experimental dawn using DAM (Fig. 4b). However, using DrosoVAM this period of activity is much less apparent, probably due to the generally higher level of nighttime activity observed.

**Figure 5.**
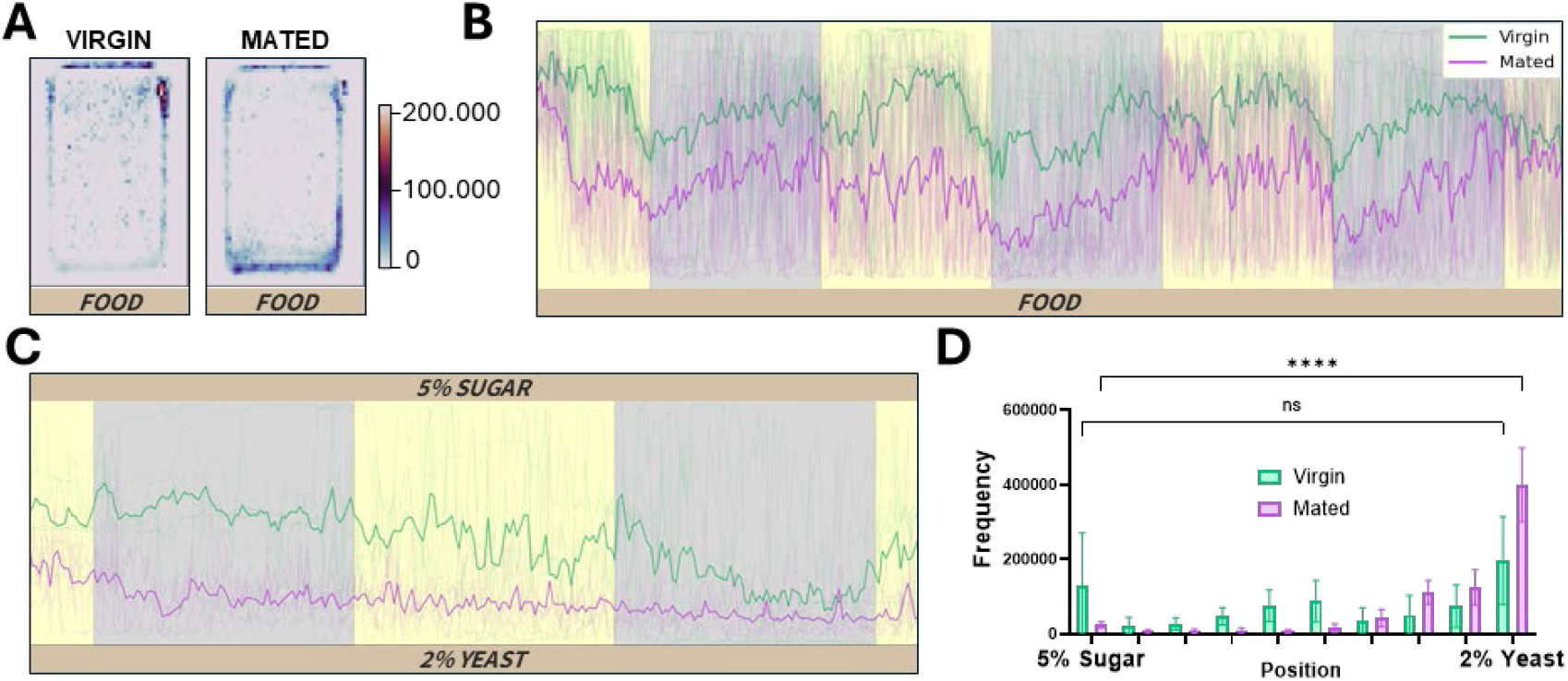
Position analysis of virgin and mated *D. melanogaster*. Virgin flies are shown in green and mated flies are shown in purple. Light and dark periods are shown with the yellow and grey background, respectively. A. Heatmaps showing the position of the flies in the chambers over a 72h period. The side of the chamber where the food is located is represented at the bottom of the heatmap. B (as indicated). Temporal representation of the position of the flies in the locomotor activity chambers. The extremity of the chamber in which food is present is located at the bottom of the chart (as indicated). Positions of all the flies are shown as thin lines, and the average position is represented with the thicker line. C. Temporal representation of the position of the flies in the food preference chambers. Sugar food is located at the top of the chart (indicated as 5% sugar), and yeast is at the bottom (indicated as 2% yeast). D. Histogram of the position of the flies along the food preference chambers. The chambers are divided in 10 bins (∼5mm per bin) and t-Test have been performed between the bins closest to the foods.

As mated females seemed to spend more time near the food, we wanted to test if DrosoVAM could be adapted to study feeding behaviors in mated females. For this, we have developed an alternative version of the monitoring chamber (Fig. S1b) that can be filled with two different types of food, allowing us to test for a possible shift in food preference. In this test, we put mated and virgin flies into chambers with 1% agar supplemented either with 2% yeast extract (source of proteins) or with 5% sucrose (source of carbohydrates). Previously, it was shown that *D. melanogaster* females change their food preference towards high protein sources after mating (Ribeiro and Dickson, 2010; Vargas et al., 2010). Demonstrating the versatility of our system, we can confirm that mated flies spent more time near the yeast containing food while virgin flies show little or no preference (Fig. 5c-d).

## Discussion

The TriKinetics DAM system has made it possible to quantify the differences in behavior between different *D. melanogaster* genotypes under different conditions (Chiu et al., 2010; Isaac et al., 2009). The system has been highly adopted due to its simplicity, and its scalability to large numbers. However, its simplification of activity to crossing an IR beam at the center of tube may lead to problems in interpreting the phenotypes being scored; movements made away from the center line are lost, while small movements near the beam can give a false sense of hyperactivity. Modern video monitoring technology has now made using such a proxy for activity unnecessary for many applications. Here, we have shown that a video-based system like DrosoVAM is able to provide comparable, if not more-complete results relative to DAM for the monitoring of *Drosophila* displacement activity. For example, previous DAM experiments have shown that mated females are more active than virgin flies during the day and described it as a post-mating reduction in siesta sleep (Isaac et al., 2009). Here, using DrosoVAM, we show that mated females actually display an almost constant level of activity that is higher than that of virgin females during periods of time in the day and night phases (Fig. 4a). Thus, the reduction in daytime siesta sleep seen in DAM, while real, is an incomplete picture of the activity changes experienced by mated females.

We can propose two likely reasons that the DAM system does not find a nighttime increase in activity. The first possibility is that the discrepancy between DAM and DrosoVAM results stems from a change in the localization of the flies that is not seen by DAM, while the second suggests that the change in behavior is linked to the difference in environment between the two systems.

Regarding the first possibility, previous work has shown that isolated females prefer to lay their eggs in the dark (Bailly et al., 2023). Thus, it seems likely that mated females would spend more time closer to the food at night to lay eggs. Our DrosoVAM results are consistent with this interpretation. While mated females are more active than virgin flies (particularly during the nighttime hours (Fig. 5b)), they tend to spend more of their time at or near the food than virgin flies. Because of this relocation, females could be more active at night, but this activity would be less likely to activate a DAM activity sensor located at the tube midline. If this interpretation is correct, it highlights an important caveat in using DAM or any system that proxies the frequency of a specific event to measure another. In many articles, periods of DAM inactivity (5 consecutive minutes without the fly crossing the central beam) have been labeled as sleep. Our work has shown that mated flies are often active on or near the food (Fig. 5a-b), potentially leading to falsely attributing egg-laying or feeding behaviors as sleep when activity is not directly monitored. While we do not discount a potential inverse proportionality of the beam-crossing behavior to sleep activity, particularly in individual males or virgin females as seen in (Cao and Edery, 2017; Dorcikova et al., 2023; Schretter et al., 2018; Shahandeh et al., 2024), our DrosoVAM experiments show that potentially unrelated changes in behavior might cause a quantitative misattribution of sleep bouts in flies.

An alternative possibility to explain the differences between the DAM and DrosoVAM results in overall activity is that flies behave differently in the confined environment of tubes relative to the more-open environment of our chambers. Indeed, in the context of DAM, the animals are only given a few millimeters of space in which they can move in two of the three axes. As the size and characteristics of the chambers in which the flies are observed have been shown to affect their behavior, this is a relevant concern (Philipsborn et al., 2023; Simon and Dickinson, 2010; Valente et al., 2007). We chose the size of our DrosoVAM chambers to allow for the monitoring of multiple flies, while providing each fly with an ample supply of food and a space sufficient for pivoting and movement in all directions. Furthermore, while a wider environment may explain the discrepancies in activity levels, it is possible that it would also affect the activity pattern itself. Indeed, in our DrosoVAM experiment on virgin females, aside from the startle activity response to light transitions, we failed to identify the stereotypical “morning anticipation”, previously characterized using narrower chambers. Additionally, studies using DAM system have found that the richness of the food may alter the sleeping behavior of mated females (Duhart et al., 2023). However, we ruled out any impact of food composition in the discrepancy we characterized, as the food used for both DAM and DrosoVAM experiments was identical.

Given the almost infinite number of variables that one can test when studying behavior, one of the strengths of DrosoVAM is that it can be easily and inexpensively modified to fit one’s needs. For example, as a follow-up to our mated female localization results, we created a second setup with two different feeding areas per chamber to monitor where mated females would choose to spend their time. Using this assay, we were able to highlight how mated females display a striking preference for rich food (2% yeast) that is not seen with virgin flies (Fig. 5 c-d). This result is consistent with previous studies using alternative methods (food dyes, etc) (Ribeiro and Dickson, 2010; Vargas et al., 2010), and shows how DrosoVAM can be quickly and easily modified to monitor different aspects of behavior without the need to invest in more expensive equipment. The subsequent analysis of DrosoVAM data can also easily be adapted “*a posteriori*”, for example to quantify sleep based on locomotor activity. While the quantification of sleep is theoretically possible, we decided to focus primarily on raw locomotor activity to assess changes in the behavior of mated females, avoiding the misattribution of sleep to immobile flies that could be eating or laying eggs. We have also been able to use DrosoVAM with other species of Drosophila like the larger flies, *D. virilis* and *D. hydei*. Preliminary results using these species with DrosoVAM actually show that the system not only works but actually functions better due to the larger size and darker IR profiles of these flies. Thus, the versatility of DrosoVAM in experiment design is not limited to modification of chambers and types of analysis, but also to the species that can be used in it.

Not only can DrosoVAM easily be implemented and modified, but it is also noticeably easier to setup an experiment compared to the DAM system. Indeed, the preparation of the chamber for monitoring can take as few as five minutes, the time necessary to pour food in the food wells. The placement of the flies in the chambers, which can take place as soon as the food is solid, also takes a limited amount of time. Altogether, setting up the experiment and starting the recording takes less than an hour allowing for the rapid start of monitoring.

Given the limitations of the DAM system, it is not surprising that several video-based monitoring systems have been developed by other groups. Many of these systems have been specifically designed to monitor more-detailed behaviors, using higher resolution cameras over shorter periods of time or with less flies to keep the data manageable (Alphen et al., 2013; Bräcker et al., 2019; Keleş et al., 2025; Niu et al., 2021; Qu et al., 2022. Still, other systems may be able to perform similar functions and at similar scales to DrosoVAM (Donelson et al., 2012; Garbe et al., 2015; Gilestro, 2012; Guo et al., 2016; Simon and Dickinson, 2010; Xu et al., 2021; Zimmerman et al., 2008). For comparison of some existing methods, see Table 1. Our choice to develop DrosoVAM was driven, not because of the lack of other systems, but because adapting these systems to our varying needs proved to be harder than developing an inexpensive, simpler, modular system using easily obtainable parts and software. Thus, DrosoVAM’s strength lies in its providing of a reliable and robust method to monitor displacement activity that is both simple to implement and to evolve as one sees fit.

**Table 1:** Comparison of video-monitoring systems.

| Reference | Number of flies | Chamber type and size | Video technology | Analysis technology | Recording duration | Studied metric |
| --- | --- | --- | --- | --- | --- | --- |
| Alphen et al., 2013 | 1 | No chamber | Webcam @3fps | pySolo, MATLAB | Several days | Micromovement and electrophysiology |
| Bräcker et al., 2019 | 1 | Food patch in 30x30x30 chamber | High speed camera | MATLAB | 1 hour | Egg Laying |
| Keleş et al., 2025 | 1 | Small ~3x~5x~7mm | High resolution camera | DeepLabCut, MATLAB, Python | 16 hours | Micromovements |
| Niu et al., 2021 | 18 x 1 | Glass tubes | Nikon Stereomicroscope @1fps | pySolo, MATLAB | 5 days | Feeding |
| Qu et al., 2022 | 72 x 1 | Flat disc of 16mm of diameter | Standard camera @30fps | Python | 3 hours | Drug testing on activity |
| Gilestro, 2012 | 32 x 1 | DAM tubes | Standard camera @~20fps | pySolo | 5 to 7 days | Sleep and locomotion monitoring |
| Simon and Dickinson, 2010 | 1 x 50 | Slopped plate, 13cm diameter | Basler camera and lens @20fps | Ctrx, Flytrax, MATLAB | 10 minutes | Locomotion (proof of concept) |
| Xu et al., 2021 | "Tens of flies" | Ellipse 30 x 30 x 3 mm | Infrared camera @25fps | MATLAB | 3 days | Locomotion |
| Guo et al., 2016 | 96 x 1 | 96-well plate | Webcam @1fps | Fiji MTrack2 plugin | 1 hour | Arousal measurement |
| Garbe et al., 2015 | 6/12/24 x 1 | 6/12/24-well plates | Standard camera @30fps | Ethovision | 24 hours | Locomotor activity |
| Donelson et al., 2012 |  | DAM tubes | Webcam @1fps | Tracker, MATLAB | 3 days | Locomotor activity |
| Zimmerman et al., 2008 | 8 x 1 | DAM tubes | Standard camera @0.2fps | MATLAB | 24 hours | Sleep |
| Revel et al., 2025 | 32 x 1 | 3D printed | Basler camera @5fps | DeepLabCut, Python | Up to 9 days | Locomotor activity |

## Supplementary figures

**Figure S1.**
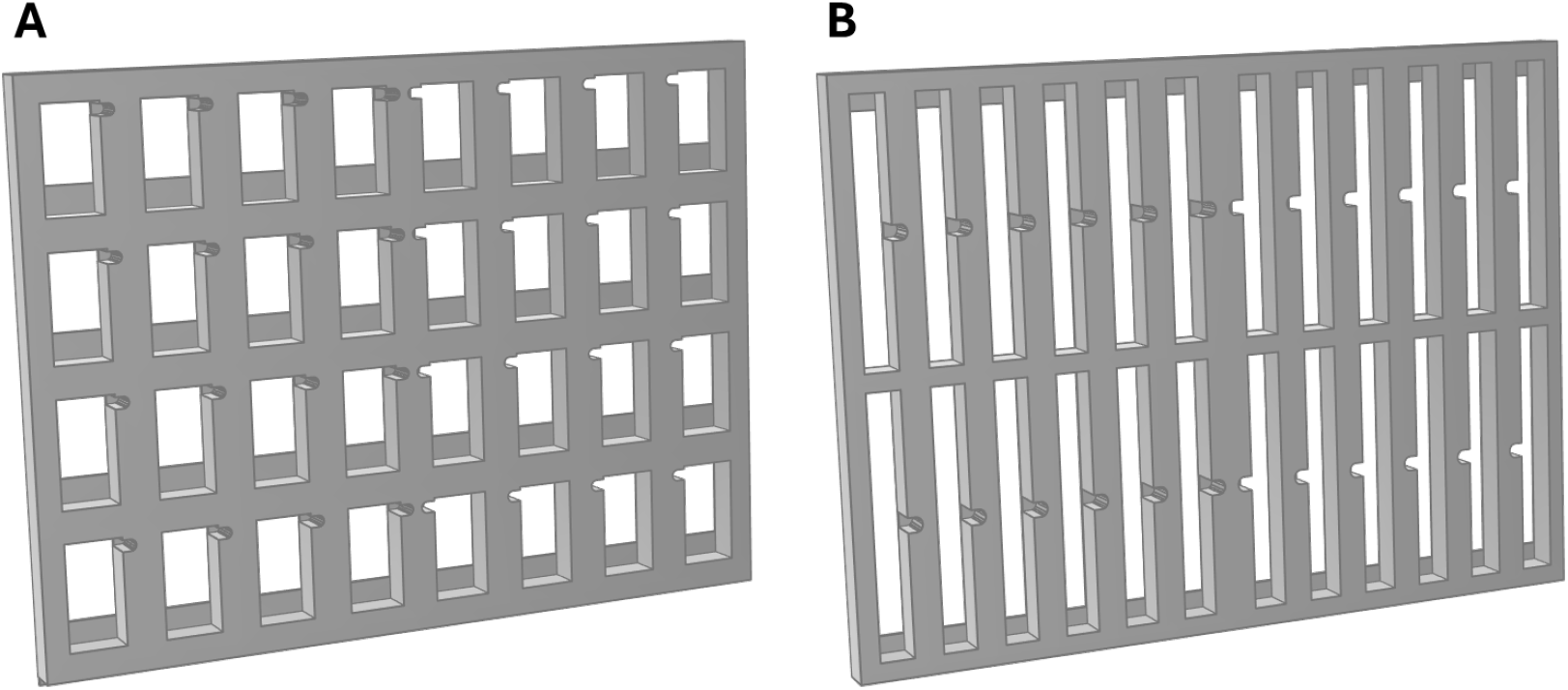
3D-printed recording arenas (196×145mm). A. Arena used for locomotor activity assays containing 32 chambers (15×25 mm), with indentations to insert the flies in the chamber and a wall support for the food at the bottom. B. Arena used for food preference assays with 24 chambers (10×50 mm). The two food supports are located at both extremities of the chambers. Chambers have holes to insert the flies using a mouth pipet (3 mm diameter).

**Figure S2.**
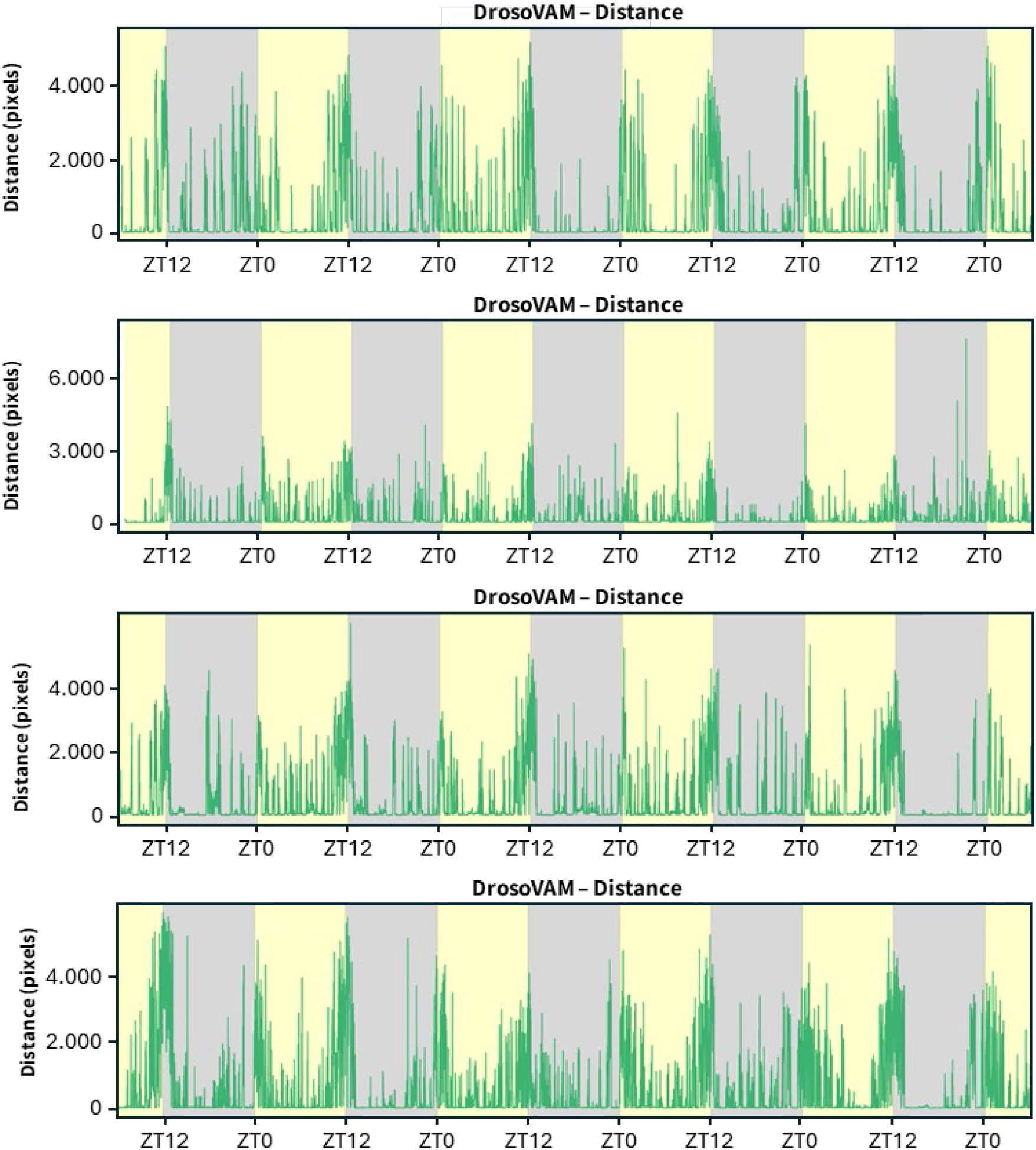
Individual locomotor activity from virgin wild-type females. Light/dark periods are shown with yellow/grey background respectively. Distance moved (in pixels) by representative individual flies in 1 minute bins estimated using DrosoVAM.

## Notes

https://github.com/LabMaeda/DrosoVAM

### Competing Interest Statement

The authors have declared no competing interest.

### Funding

- Swiss National Science Foundation: 310030_192621, 320030_227485

## References

Alphen, B. van, Yap, M.H.W., Kirszenblat, L., Kottler, B. Swinderen, B. van, 2013. A Dynamic Deep Sleep Stage in Drosophila. J. Neurosci. 33, 6917–6927. 10.1523/jneurosci.0061-13.2013

Bailly, T.P.M., Kohlmeier, P., Etienne, R.S., Wertheim, B., Billeter, J.-C., 2023. Social modulation of oogenesis and egg laying in Drosophila melanogaster. Curr. Biol. 33, 2865–2877.e4.

Bräcker, L.B., Schmid, C.A., Bolini, V.A., Holz, C.A., Prud’homme, B., Sirota, A., Gompel, N., 2019. Quantitative and Discrete Evolutionary Changes in the Egg-Laying Behavior of Single Drosophila Females. Front. Behav. Neurosci. 13, 118.

Brenman‐Suttner, D.B., Yost, R.T., Frame, A.K., Robinson, J.W., Moehring, A.J., Simon, A.F., 2020. Social behavior and aging: A fly model. Genes, Brain Behav. 19, e12598. 10.1111/gbb.12598

Busto, G.U., Guven-Ozkan, T., Fulga, T.A., Vactor, D.V., Davis, R.L., 2015. microRNAs That Promote or Inhibit Memory Formation in Drosophila melanogaster. Genetics 200, 569–580.

Cao, W., Edery, I., 2017. Mid-day siesta in natural populations of D. melanogaster from Africa exhibits an altitudinal cline and is regulated by splicing of a thermosensitive intron in the period clock gene. BMC Evol. Biol. 17, 32.

Chiu, J.C., Low, K.H., Pike, D.H., Yildirim, E., Edery, I., 2010. Assaying Locomotor Activity to Study Circadian Rhythms and Sleep Parameters in Drosophila. J. Vis. Exp. : JoVE 2157.

Cooper, J.F., Raamsdonk, J.M.V., 2018. Modeling Parkinson’s Disease in C. elegans. J. Park.’s Dis. 8, 17–32. 10.3233/jpd-171258

Donelson, N.C., Donelson, N., Kim, E.Z., Slawson, J.B., Vecsey, C.G., Huber, R., Griffith, L.C., 2012. High-Resolution Positional Tracking for Long-Term Analysis of Drosophila Sleep and Locomotion Using the “Tracker” Program. PLoS ONE 7, e37250. 10.1371/journal.pone.0037250

Dorcikova, M.M., Duret, L.C., Pottié, E., Nagoshi, E., 2023. Circadian clock disruption promotes the degeneration of dopaminergic neurons in male Drosophila. Nat. Commun. 14, 5908.

Dubowy, C., Sehgal, A., 2017. Circadian Rhythms and Sleep in Drosophila melanogaster. Genetics 205, 1373–1397.

Duhart, J.M., Buchler, J.R., Inami, S., Kennedy, K.J., Jenny, B.P., Afonso, D.J.S., Koh, K., 2023. Modulation and neural correlates of postmating sleep plasticity in Drosophila females. Curr. Biol. 33, 2702–2716.e3. 10.1016/j.cub.2023.05.054

Frank, K.D., Zimmerman, W.F., 1969. Action Spectra for Phase Shifts of a Circadian Rhythm in Drosophila. Science 163, 688–689.

Garbe, D.S., Bollinger, W.L., Vigderman, A., Masek, P., Gertowski, J., Sehgal, A., Keene, A.C., 2015. Context-specific comparison of sleep acquisition systems in Drosophila. Biol. Open 4, 1558–1568. 10.1242/bio.013011

Gaspar, M., Dias, S., Vasconcelos, M.L., 2022. Mating pair drives aggressive behavior in female Drosophila. Curr. Biol. 32, 4734–4742.e4.

Gilestro, G.F., 2012. Video tracking and analysis of sleep in Drosophila melanogaster. Nat. Protoc. 7, 995–1007.

Guo, F., Yu, J., Jung, H.J., Abruzzi, K.C., Luo, W., Griffith, L.C., Rosbash, M., 2016. Circadian neuron feedback controls the Drosophila sleep–activity profile. Nature 536, 292–297. 10.1038/nature19097

Isaac, R.E., Li, C., Leedale, A.E., Shirras, A.D., 2009. Drosophila male sex peptide inhibits siesta sleep and promotes locomotor activity in the post-mated female. Proc. R. Soc. B: Biol. Sci. 277, 65–70.

Keleş, M.F., Sapci, A.O.B., Brody, C., Palmer, I., Mehta, A., Ahmadi, S., Le, C., Taştan, Ö., Keleş, S., Wu, M.N., 2025. FlyVISTA, an integrated machine learning platform for deep phenotyping of sleep in Drosophila. Sci. Adv. 11, eadq8131.

Konopka, R.J., Benzer, S., 1971. Clock Mutants of Drosophila melanogaster. Proc. Natl. Acad. Sci. 68, 2112–2116. 10.1073/pnas.68.9.2112

Martín, F., Alcorta, E., 2017. Novel genetic approaches to behavior in Drosophila. J. Neurogenet. 31, 288–299. 10.1080/01677063.2017.1395875

Mathis, A., Mamidanna, P., Cury, K.M., Abe, T., Murthy, V.N., Mathis, M.W., Bethge, M., 2018. DeepLabCut: markerless pose estimation of user-defined body parts with deep learning. Nat. Neurosci. 21, 1281–1289.

Mirth, C.K., Alves, A.N., Piper, M.D., 2019. Turning food into eggs: insights from nutritional biology and developmental physiology of Drosophila. Curr. Opin. Insect Sci. 31, 49–57.

Nagoshi, E., 2018. Drosophila Models of Sporadic Parkinson’s Disease. Int. J. Mol. Sci. 19, 3343.

Nath, T., Mathis, A., Chen, A.C., Patel, A., Bethge, M., Mathis, M.W., 2019. Using DeepLabCut for 3D markerless pose estimation across species and behaviors. Nat. Protoc. 14, 2152–2176.

Niu, M., Zhang, X., Li, W., Wang, J., Li, Y., 2021. dFRAME: A Video Recording-Based Analytical Method for Studying Feeding Rhythm in Drosophila. Front. Genet. 12, 763200.

Orger, M.B., Polavieja G.G. de, 2015. Zebrafish Behavior: Opportunities and Challenges. Annu. Rev. Neurosci. 40, 1–23. 10.1146/annurev-neuro-071714-033857

Philipsborn, A.C. von, Liu, T., Yu, J.Y., Masser, C., Bidaye, S.S., Dickson, B.J., 2011. Neuronal Control of Drosophila Courtship Song. Neuron 69, 509–522.

Philipsborn, A.C. von, Shohat-Ophir, G., Rezaval, C., 2023. Single-Pair Courtship and Competition Assays in Drosophila. Cold Spring Harb. Protoc. 2023, pdb.prot108105.

Qu, S., Zhu, Q., Zhou, H., Gao, Y., Wei, Y., Ma, Y., Wang, Z., Sun, X., Zhang, Lei, Yang, Q., Kong, L., Zhang, Li, 2022. EasyFlyTracker: A Simple Video Tracking Python Package for Analyzing Adult Drosophila Locomotor and Sleep Activity to Facilitate Revealing the Effect of Psychiatric Drugs. Front. Behav. Neurosci. 15, 809665.

Raun, N., Jones, S., Kramer, J.M., 2021. Conditioned courtship suppression in Drosophila melanogaster. J. Neurogenet. 35, 154–167.

Ribeiro, C., Dickson, B.J., 2010. Sex Peptide Receptor and Neuronal TOR/S6K Signaling Modulate Nutrient Balancing in Drosophila. Curr. Biol. 20, 1000–1005.

Riva, S., Ispizua, J.I., Breide, M.T., Polcowñuk, S., Lobera, J.R., Ceriani, M.F., Risau-Gusman, S., Franco, D.L., 2022. Mating disrupts morning anticipation in Drosophila melanogaster females. Plos Genet 18, e1010258.

Schretter, C.E., Vielmetter, J., Bartos, I., Marka, Z., Marka, S., Argade, S., Mazmanian, S.K., 2018. A gut microbial factor modulates locomotor behaviour in Drosophila. Nature 563, 402–406.

Shahandeh, M.P., Abuin, L., Decker, L.L.D., Cergneux, J., Koch, R., Nagoshi, E., Benton, R., 2024. Circadian plasticity evolves through regulatory changes in a neuropeptide gene. Nature 635, 951–959.

Simon, J.C., Dickinson, M.H., 2010. A New Chamber for Studying the Behavior of Drosophila. PLoS ONE 5, e8793.

Sitaraman, D., Vecsey, C.G., Koochagian, C., 2024. Activity Monitoring for Analysis of Sleep in Drosophila melanogaster. Cold Spring Harb. Protoc. 2024, pdb.top108095. 10.1101/pdb.top108095

Tomioka, K., Sakamoto, M., Harui, Y., Matsumoto, N., Matsumoto, A., 1998. Light and temperature cooperate to regulate the circadian locomotor rhythm of wild type and period mutants of Drosophila melanogaster. J. Insect Physiol. 44, 587–596.

Tsuda, L., Lim, Y.-M., 2018. Drosophila Models for Human Diseases. Adv. Exp. Med. Biol. 1076, 25–40.

Tully, T., Preat, T., Boynton, S.C., Vecchio, M.D., 1994. Genetic dissection of consolidated memory in Drosophila. Cell 79, 35–47.

Valente, D., Golani, I., Mitra, P.P., 2007. Analysis of the Trajectory of Drosophila melanogaster in a Circular Open Field Arena. PLoS ONE 2, e1083.

Vargas, M.A., Luo, N., Yamaguchi, A., Kapahi, P., 2010. A Role for S6 Kinase and Serotonin in Postmating Dietary Switch and Balance of Nutrients in D. melanogaster. Curr. Biol. 20, 1006–1011.

White, B.H., 2016. What genetic model organisms offer the study of behavior and neural circuits. J. Neurogenet. 30, 54–61. 10.1080/01677063.2016.1177049

Wu, M.N., 2025. FlyVISTA, an integrated machine learning platform for deep phenotyping of sleep in Drosophila. Sci. Adv. 11, eadq8131. 10.1126/sciadv.adq8131

Xu, X., Yang, W., Tian, B., Sui, X., Chi, W., Rao, Y., Tang, C., 2021. Quantitative investigation reveals distinct phases in Drosophila sleep. Commun. Biol. 4, 364.

Yamamoto, D., Koganezawa, M., 2013. Genes and circuits of courtship behaviour in Drosophila males. Nat. Rev. Neurosci. 14, 681–692.

Zimmerman, J.E., Raizen, D.M., Maycock, M.H., Maislin, G., Pack, A.I., 2008. A Video Method to Study Drosophila Sleep. Sleep 31, 1587–1600. 10.1093/sleep/31.11.1587

Zwarts, L., Versteven, M., Callaerts, P., 2012. Genetics and neurobiology of aggression in Drosophila. Fly 6, 35–48.

